# Protein degradation analysis by affinity microfluidics

**DOI:** 10.1101/2021.10.13.464189

**Authors:** Lev Brio, Danit Wasserman, Efrat Michaely-Barbiro, Doron Gerber, Amit Tzur

## Abstract

Protein degradation mediated by the ubiquitin-proteasome pathway regulates signaling events in all eukaryotic cells, with implications in pathological conditions such as cancer and neurodegenerative diseases. Detection of protein degradation is an elementary need in basic and translational research. In vitro degradation assays, in particular, have been instrumental in the understanding of how cell proliferation and other fundamental cellular processes are regulated. These assays are direct, quantitative and highly informative but also laborious, typically relying on low-throughput polyacrylamide gel-electrophoresis followed by autoradiography or immunoblotting. We present protein degradation on chip (pDOC), a MITOMI-based integrated microfluidic device for discovery and analysis of ubiquitin-mediated proteolysis. The platform accommodates microchambers on which protein degradation is assayed quickly and simultaneously in physiologically relevant environments, using minute amount of reagents. Essentially, pDOC provides a multiplexed, sensitive and colorimetric alternative to the conventional degradation assays, with relevance to biomedical and translational research.

## Introduction

Protein degradation by the ubiquitin-proteasome system is a central regulatory module through which the level of proteins in all eukaryotic cells remains balanced. Deviation from the desired amount of each protein at any given moment can be detrimental to the cell, leading to dysfunctional tissues and a wide range of illnesses in human, including cancer, cystic fibrosis, and neurodegenerative diseases [1] [2].

The core cascade underlying ubiquitination involves three enzymes: The E1 enzyme covalently binds and activates the ubiquitin molecule for transfer to an E2 conjugating enzyme. Then, the ubiquitin-conjugated E2 interacts with an E3 ubiquitin-ligase enzyme, which catalyzes the transfer of ubiquitin molecules from the E2 to the target protein or to a second ubiquitin molecule, typically via an isopeptide bond to a lysine residue. Finally, a target protein that is covalently bound to a chain of ubiquitin moieties can be recognized by the proteasome for degradation [1, 2]. Hundreds of different E3 enzymes underlie the enormous functional reach and specificity of the entire ubiquitination process. With respect to cell proliferation and cell cycle regulation, the ubiquitin ligases <u>a</u>naphase-<u>p</u>romoting <u>c</u>omplex/<u>c</u>yclosome (APC/C) and <u>S</u>kp1-<u>C</u>ullin-<u>F</u>-box protein complex (SCF) are particularly important [3–5]. The substrate specificity of both complexes is dependent on coactivators: Cdc2O and Cdh1 for the APC/C and one of several F-box proteins for the SCF, e.g., Skp2 and β-TrCP [6–8]. Overall, orderly proteolysis mediated by cell cycle regulated E3 enzymes ensures unidirectional cell cycle in all eukaryotes. [6–10].

Protein degradation, however, cannot be automatically inferred from ubiquitination; while some forms of ubiquitin chains trigger proteolysis, monoubiquitination and other forms of polyubiquitination regulate signaling cascades via proteasome-independent pathways [11]. Furthermore, ubiquitination can be reversed by enzymes called deubiquitinases in a manner that can prevent proteolysis [12]. On the flip side, proteasomal degradation may not always be coupled to ubiquitination [13]. Thus, protein degradation must be determined directly.

Protein degradation assays in cell-free extracts, also known as ‘cell-free systems’, have been instrumental in cell biology research, enabling direct and quantitative analyses of ubiquitin-mediated proteolysis in physiologically relevant environments. In fact, much of the cell cycle principles were discovered by monitoring the degradation of cell cycle proteins in extracts from frog eggs or cycling human cells (see for example [14–20]). The extensive use of these ‘degradation assays’ in today’s modern era, is a testament to their efficacy (see for example [21–23]). Interestingly, conventional degradation assays have never truly benefited from modern technologies, and still typically rely on gel-electrophoresis, autoradiography and large amount of biological material.

Integrated microfluidics and pneumatic microvalves paved the way to protein chips in which the arrayed proteins are freshly expressed in a physiological environment that maintain proper protein folding and activity [24]. The target proteins are expressed either on chip or externally, and subsequently immobilized to microchambers via a designated surface chemistry. Then, a large panel of direct colorimetric assays can be performed over thousands of microchambers, using minute amounts of reagents. In recent years, we developed several microfluidic devices based on <u>m</u>echanically <u>i</u>nduced <u>t</u>rapping <u>o</u>f <u>m</u>olecular <u>i</u>nteractions (MITOMI), with which we discovered and detected i) protein interactions with DNA, RNA, proteins and viruses; and ii) protein post-translational modification (PTM), specifically, phosphorylation, autophosphorylation and ubiquitination [24–30].

The combination of integrated microfluidics, protein arrays and cell-free systems from healthy or pathological sources holds great potential in biomedical research and diagnostics. In this study, we utilize the MITOMI platform for protein degradation analyses. The proof of concept is demonstrated using cell extracts with APC/C-specific activity. The method, named pDOC (<u>p</u>rotein <u>d</u>egradation <u>o</u>n <u>c</u>hip) provides a fast, sensitive and costeffective alternative to the classic method by which proteasome-mediated proteolysis has been assayed in vitro almost unvaryingly for nearly half a century.

## Materials and methods

### Plasmids

pCS2-Flag-FA vector was generated by annealing Flag tag oligos and ligating final fragment into pCS2-FA vector using BamHI and FseI restriction sites. pCS2-Flag-FA-Securin-GFP w.t and Δ64 variant plasmids were generated by cloning w.t or Δ64 Securin-GFP [27] into pCS2-Flag-FA vector, using FseI (5’) and AscI (3’) flanked primers. pCS2-GemininΔ27-GFP was generated by deleting amino acids 1-27 from Geminin-GFP [27] using QuikChange^®^ Lightning mutagenesis kit (Agilent, 210513). The plasmids pCS2-Flag-FA-Geminin-GFP w.t and Δ27 variant were generated by cloning Geminin-GFP into pCS2-Flag-FA vector sing FseI and AscI restriction sites. pCS2-Flag-FA-p27-GFP was generated by replacing Geminin open reading frame (ORF) with p27 ORF using FseI and AgeI restriction sites and pCS2-FA-p27 as a template [27]. Flag-p27-myc fragment was generated by a two-step assembly PCR using pCS2-Flag-FA-p27-GFP template, a first primer set containing a c-Flag tag (5’) and a Myc tag (3’), and a second primer set containing a T7 promoter (5’) and a T7 terminator sequence (3’). All GFP-tagged proteins caried enhanced variant of GFP (eGFP).

### Cell culture maintenance

NDB cells are based on the HEK293 cell line. A detailed description of this cell system can be found in Ref [18]. NDB and HeLa S3 (ATCC; #CCL-2.2) cells were maintained in tissue culture dishes containing Dulbecco’s Modified Eagles Medium (DMEM) supplemented with 10% fetal bovine serum, 2 mM L-glutamine, and 1% Penicillin-Streptomycin solution (Biological Industries; #01-055-1A, #04-001-1A, #03-020-1B, #03-031-1B). Cells were maintained at 37°C in a humidified 5% CO2-containing atmosphere. HeLa S3 cells were either cultured on dishes or in 1-l glass spinner flasks in suspension (80 rpm). NDB cells were cultured in the presence of 5μg/ml Blasticidin (Life Technologies; #A11139-03) to maintain the pcDNA6/TR plasmid carrying the ORF for Tet repressor.

### Cell synchronization

For late-mitosis synchronization, NDB cells were cultured in 150 mm/diameter dishes. After reaching a confluency of about 75%, the cells were treated with 1 μg/ml Tetracycline (Sigma-Aldrich; #87128) for 22 hr and harvest for extract preparation. For S-phase synchronization, HeLa S3 cells were cultured in suspension for 72 h up to a concentration of approximately 5×105 cells/ml. Cells were then supplemented with 2 mM Thymidine for 22 hr, washed with DMEM (twice, 5 min, 250×g) and released into prewarmed fresh media (37°C) for additional 9 hr. Cell culture was then supplemented again with 2 mM Thymidine for 19 h before harvest for extract preparation.

### Preparation of cell extracts

HeLa S3 extracts: S-phase Synchronous HeLa S3 cells were washed with ice-cold 1× PBS and lysed in a swelling buffer (20 mM HEPES, pH 7.5, 2 mM MgCl_2_, 5 mM KCl, 1 mM Dithiothreitol [DTT], and protease inhibitor cocktail [Roche; #11836170001]) supplemented with energy-regenerating mixture, E-mix (1 mM ATP, 0.1 mM ethylene glycol-bis [β-aminoethyl ether]-N,N,N’,N’-tetra acetic acid [EGTA], 1 mM MgCl_2_, 7.5 mM creatine phosphate, 50 μg/ml creatine phosphokinase). Cells were incubated on ice for 30 min and homogenized by freeze–thawing cycles in liquid nitrogen and passed through a 21-G needle for 10 times. Extracts were cleared by subsequent centrifugation (17,000 × g; 10 and 40 min), and stored at −80°C. NDB mitotic extracts: Tet-induced NDB cells were collected from 20-24 150 mm dishes by gentle wash with ice-cold PBS. Extracts were prepared as described for HeLa S3. For more details see [18][31].

### In vitro expression of target proteins

Target proteins were *in-vitro* expressed using rabbit reticulocyte lysate (TNT-coupled reticulocyte system; Promega; #L4600, #L4610) supplemented with either ^35^S-methionine/S-L-cysteine mix (PerkinElmer; #NEG772002MC) for radiography detection or with untagged Methionine (Promega #L118A) and Green Lysine (FluoroTect™ GreenLys, Promega #L5001).

### Off-chip Degradation Assay

Degradation assays were performed in 20 μl cell extract supplemented with 1 μl of 20× energy regenerating mixture (see above), 1 μl of 10 mg/ml Ub solution (Boston Biochem; #U-100H), and 1μl radiolabeled in vitro translated protein of interest. For a negative control, reaction mixture was supplemented with proteasome inhibitor MG132 (20 μM; Boston Biochem; #I-130). Reaction mixtures were incubated at 28°C, and samples of 4-5 μl were collected in 15-20 min intervals. **<u>Off-chip detection</u>**: Time-point samples were mixed with 4× Laemmli Sample Buffer (BIO-RAD #1610747), denaturized (10 min, 95°C), and resolved by SDS-PAGE. Gels were soaked in a Methanol/Acetic acid (10/7.5%) fixative solution for 20 min, dried in vacuum and heat, and exposed to phosphor screen (Fuji) for 24-72 hr. In vitro translated proteins were visualized by autoradiography using Typhoon FLA 9500 Phosphorimager (GE Healthcare Life Sciences). Signal intensity (corrected for background signal) was measured by ImageJ software and was normalized to the signal at *t*_0_. All plots were created using Microsoft Excel software, version 16.20. Mean and SE values were calculated from three or four independent degradation assays. **<u>On-chip detection</u>**: Time-point samples were immediately frozen in liquid nitrogen. Before detection, samples were thawed on ice, flown through the chip for 3-5 min, and immobilized to protein chambers under the ‘button’ valve (see ‘Surface chemistry’ below). Next, the ‘button’ valves were closed, allowing unbound material to be washed by PBS. The level of target proteins (before and after degradation reactions) were determined by 488 nm-excitation and an 535/25 nm emission filter. Protein level could also be measured by immunofluorescence using fluorescently labeled antibodies (anti-Flag-Alexa 647, #15009; Cell Signaling, Danvers, MA, USA). These antibodies were flowed into the device and incubated with the immobilized proteins under the ‘button’ for 20 min at RT. Unbound antibodies were mechanically washed by PBS following the closing of the ‘button’ valve. Here, target protein levels were determined by 633 nm-excitation and an 692/40 nm emission filter.

### Device fabrication

The microfluidic device is made of two layers of PDMS. The silicon wafers are written by photolithography (Heidelberg MLA 150). Then after, the soft lithography phase is induced using silicon elastomer polydimethylsiloxane (PDMS, SYLGARD 184, Dow Corning, USA) and its curing agent to fabricate the microfluidic devices. The microfluidic devices are consisting of two aligned PDMS layers, the flow and the control layers which are prepared using different ratios of PDMS and its curing agent; 5:1 and 20:1 for the control and flow layers, respectively. The control layer is degassed and baked for 30 min at 80°C. The flow layer is initially spin coated (Laurell, USA) at 2000 rpm for 60 sec and baked at 80°C for 30 min. Next, the flow and control layers are aligned using an automatic aligner machine (custom made) under a stereoscope and baked for 1.5h at 80°C for final bonding. The two-layer device is then peeled off from the wafer and bound to a cover slip glass via plasma treatment (air, 30%, 30 sec).

### Surface chemistry

Biotinylated-BSA (1 μg/μl, Thermo) is flowed for 25 min through the device, allowing its binding to the epoxy surface. On top of the biotinylated-BSA, 0.5 μg/μl of Neutravidin (Pierce, Rockford, IL) is added (flow for 20 min). The ‘button’ valve is then closed, and biotinylated-PEG (1 μg/μl, (PG2-AMBN-5k, Nanocs Inc.) is flowed over for 20 min, passivating the flow layer, except for the buttons area. Following passivation, the ‘button’ valve is released and a flow of 0.2 μg/μl biotinylated anti-GFP antibodies (Abcam; #ab6658, Cambridge, United Kingdom) or 0.01 μg/μl biotinylated anti-Flag antibodies (Cell Signaling; #2908S Danvers, MA, USA) were applied. The antibodies bound to the exposed Neutravidin, specifically to the area under the ‘button’, creating an array of anti-GFP – or anti-Flag tag. PBS buffer was used for washing in between steps. In the case of p27 immobilization, surface chemistry was performed with 0.2 μg/ml donkey anti-mouse whole IgG antibodies (#715-065-150, Jackson Immuno research laboratories, Maryland, USA) followed by 20 min flow of 6.5 μg/ml anti p27 antibodies (Santa Cruz biotechnology, Heidelberg Germany; #1641 mouse).

### On-chip degradation assay

Flag-Securin-GFP (w.t and Δ64 mut) and p27-GFP IVT products were flowed into the chip and immobilized on the surface under the ‘button’ at the protein chambers via its GFP tag, following by PBS buffer wash and scanned. Next, the ‘button’ valves were opened and the extract reaction mixtures were incubated with the protein chambers for 60 min (30°C). During the reaction, the level of the remining target protein was determined by GFP signal every 15 min. The decline in GFP signal correlated with degradation. After background signal subtraction, GFP signals were normalized to the signal at *t*_0_ (value of 1) or between 1 (max signal) and 0 (min signal).

### Image and data analysis

LS reloaded microarray scanner, GenePix7.0 (Molecular Devices) and ImageJ image analysis software were used for analysis and presentation of the images. The signal measured around the button valve was considered as the background, since no immobilization of proteins was expected there. Yet, some background signal is always detected, which results from non-specific attachment of antibodies to the device surface. We subtracted the background signal around the buttons in a ring the size of 2R with 2-pixel spacing (see supplementary material in [27]).

### Immunoblotting

Protein samples were mixed with x4 Laemmli buffer, denatured (10 min, 96°C), and resolved on freshly made 10% acrylamide gel using a Tris-glycine running buffer. Proteins were then electro-transferred onto a nitrocellulose membrane (Bio-Rad; #162-0115) using Trans-Blot Turbo transfer system (Bio-Rad). Ponceau S Solution (Sigma-Aldrich; #81462) was used to verify transfer quality. Membrane was washed (TBS), blocked (5% skimmed milk in TBST), and incubated (RT, 1 hr) with antibody solution (2.5% BSA and 0.05% sodium azide in PBS) before blotted with anti-Securin (Abcam; #AB3305) primary antibody (RT, 2 hrs). Anti-mouse Horseradish peroxidase-conjugated secondary antibody was purchased from Jackson ImmunoResearch (#115-035-003). ECL Signal was detected using EZ-ECL (Biological Industries; #20-500-171).

## Results

pDOC is based on a MITOMI device, an integrated microfluidic chip originally developed to quantify protein-ligand interactions at equilibrium [24, 32]. The basic design was modified to contain an array of 32 by 32 microcompartments. Each compartment is separated into two chambers and controlled by three valves: a ‘neck’ valve that controls the diffusion (mixing) of material from chamber I into the ‘Protein chamber’, in which a specific target protein is trapped; a ‘sandwich’ valve that separates between cell unites; and the MITOMI ‘button’ valve, which traps interacting molecules beneath it, thus taking a snapshot of the interaction at equilibrium (Figure 1A). The pDOC chip design includes separation of the master control of the three valves into sections of the chip that can be activated and controlled independently. Compared to the control of each valve type for the entire chip, this separation not only improves valve response, but more importantly, it permits time-response assays on chip. Within each section, experiments are performed in all cell units in parallel, enabling high-throughput applications of the kind shown in our previous devices [24–26, 33].

**Figure 1:**
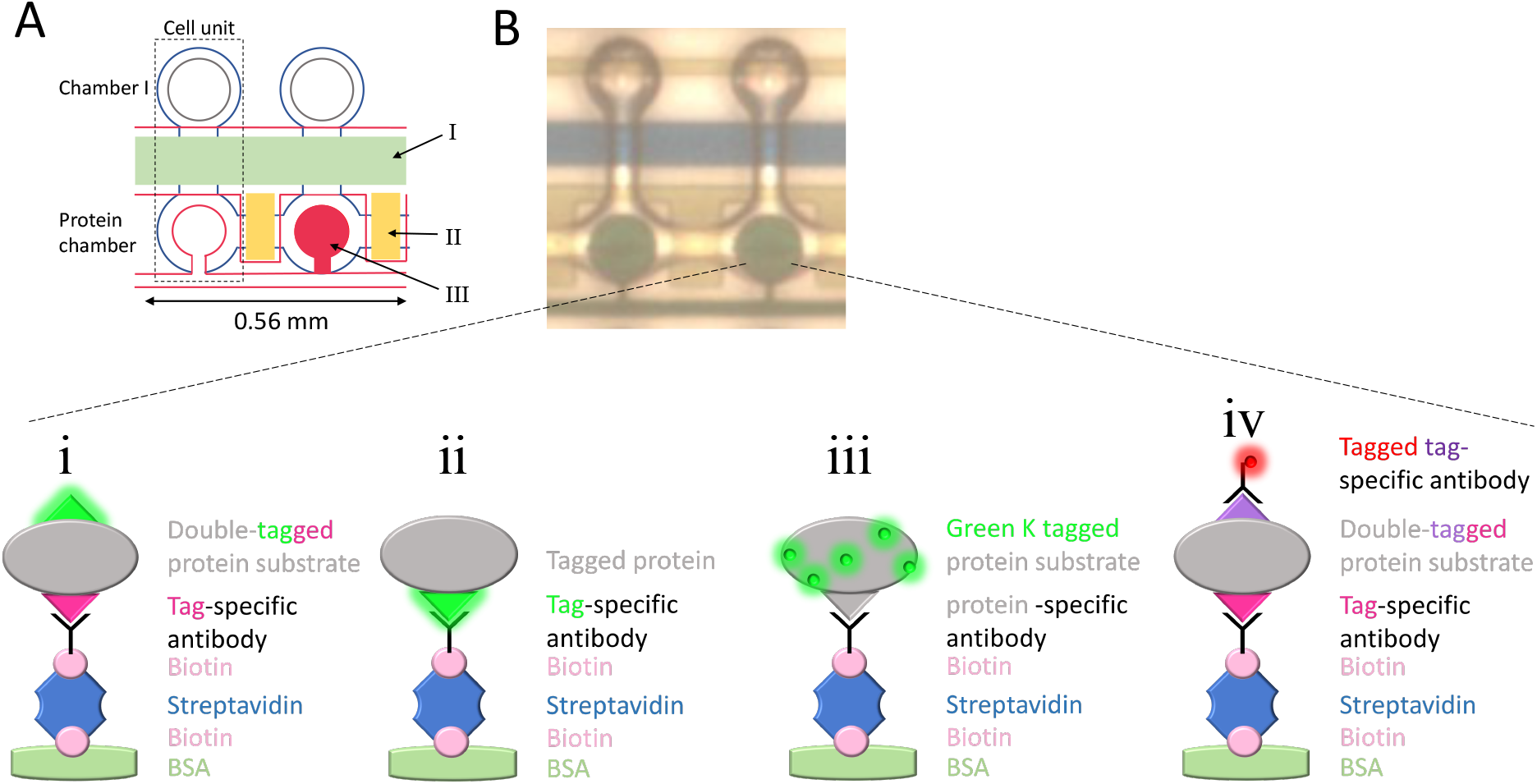
The pDOC device, surface chemistry and detection of target proteins. **(A)** An illustration of two cell units within MITOMI-based chip. Each cell unit (see black frame) comprises two chambers controlled by three pneumatic integrated valves. The two chambers are separated by the ‘neck valve’ (I). Different cell units are separated via sandwich valves (II). Samples containing target proteins (IVT products) are loaded into the protein chamber and immobilized via biotinylated antibodies that can be either protein- or tag specific. The target proteins are trapped via the MITOMI button valve (III) for quantification while the remaining unbound biomaterials are washed away. **(B)** Target proteins can be immobilized and quantified in several ways (illustrated): i) An example of a target protein carrying a fluorescent tag (e.g., GFP) for detection (see green glowing tag) and a non-fluorescent tag for immobilization; ii) A target protein tagged with GFP for both immobilization (by anti-GFP antibodies) and detection; iii) The target protein is untagged. Detection is based on fluorescent Lysine (Lys) incorporated during in vitro translation. Immobilization is via protein-specific antibodies. iv) The target protein is immobilized via tag- or protein-specific antibodies and detected by immunofluorescence via tag-specific antibodies coupled to fluorophore. Overall, on-chip immobilization of target proteins relies on biotinylated antibodies. Immobilization via non-biotinylated antibodies is possible if surface chemistry includes biotinylated IgG.

Similar to the classic degradation assays, target proteins for pDOC analyses are in vitro translated (IVT) using rabbit reticulocyte lysate that allows correct folding and PTM. However, quantitative detection is based on fluorescence rather than radioactivity. The IVT products are immobilized on the glass surface of the chip via biotin-avidin binding. To this end, specific biotinylated antibodies are applied under the button valve. Following pull down of the target protein the unbound material, such as reticulocyte lysate and cell extract, is washed away. Then the entire chip is passivated by PEG-Biotin, except for the area beneath the button (Figure. 1A; for more details, see our previous publications [25, 26, 33]. The freedom to control flow in individual sections enables multiple regimes of surface chemistry on one chip.

The device is compatible with multiple strategies of surface chemistry and possible experimental setups (illustrated in Figure 1B): i) The target protein is tagged on both N’ and C’ termini. One tag is used for immobilization via tag-specific biotinylated antibodies and the second tag is green fluorescent protein (GFP), which is used for detection. ii) The target protein is single-tagged with GFP, which is used for both immobilization and detection. iii) The target protein is double tagged. Here, however, detection is based on fluorescently labeled antibodies against short non-fluorescent tags (e.g., Flag). iv) The target protein is in vitro translated in lysate containing fluorescent lysine and immobilized by protein-specific antibodies. Importantly, immobilization can be performed with non-biotinylated antibodies if the surface chemistry also includes biotinylated IgG. Overall, the flexibility of the method simplifies assay optimization according to specific needs and limitations.

### pDOC facilitates analysis of protein degradation in cell-free extracts

Conceptually, analysis of protein degradation by pDOC is direct, simple and fast; signal detection is based on in situ quantification of fluorescent signals, thus obviating gel-electrophoresis and any other gel-related procedures, e.g fixation, drying, autoradiography or immunoblotting, and long exposures. pDOC functions like a chromatography column; it isolates and concentrates the target protein on the protein chamber of each cell unit. Our first goal was to examine whether the signal sensitivity and dynamic range of pDOC enables time-based quantification of IVT products following incubation with cell extracts in tube. As a proof of concept, we utilized mitotic extracts from HEK293 cells that are blocked in an anaphase-like state due to high levels of <u>n</u>on-<u>d</u>egradable Cyclin-<u>B</u>1. This mitotic cell-free system, hereafter referred to as NDB, recapitulates APC/C^Cdc20^-mediated proteolysis of the cell cycle proteins Securin and Geminin [18, 31].

Conventional degradation assays of radiolabeled Flag-Securin-GFP and Flag-Geminin-GFP (IVT products) in NDB mitotic extracts are shown in Figure 2A. Control experiments with non-degradable mutant variants (Geminin Δ27 and Securin Δ64) demonstrate the specificity of the assay. Equivalent experiments were performed with non-radioactive IVT products. First, we performed an end-point assay. After 60 min incubation, reaction samples were loaded on pDOC through separate channels and scanned for GFP fluorescence. Control reactions, in which IVT products were incubated in PBS, allowed us to normalize the level of each target protein at *t*_60 min_ (extracts/PBS ratio) and to estimate background signals. Overall, on-chip detection demonstrates a sharp reduction in the level of Geminin and Securin following incubation in NDB mitotic extracts, whereas non-degradable variants remained stable, exhibiting ~80% of the control GFP signals in PBS. At this juncture, we noted that background signals from reticulocyte lysate, cell extracts, and non-specific immobilization were minor (Figure S1).

**Figure 2:**
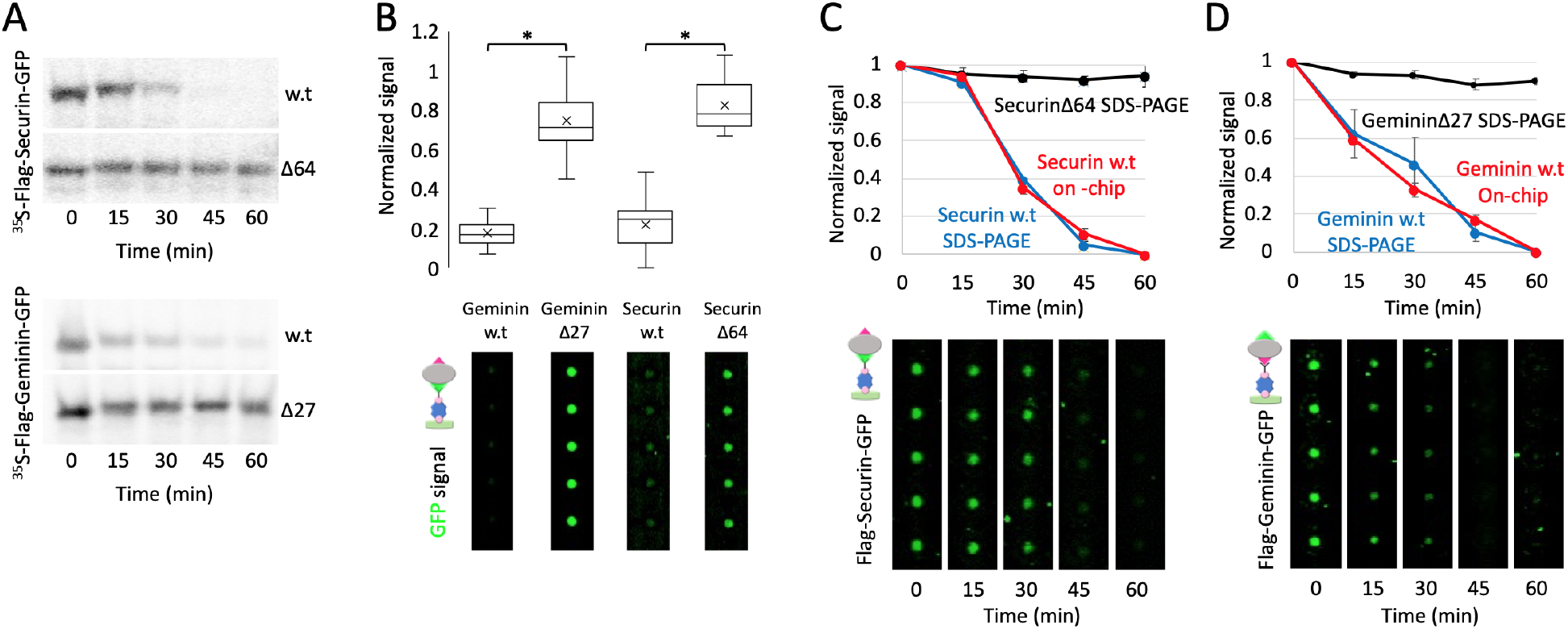
On-chip analysis of protein degradation assays. **(A)** Time-dependent degradation of ^35^S-labeled Flag-Securin-GFP, Flag-Geminin-GFP and their non-degradable variants (Δ64 and Δ27, respectively) in NDB mitotic extracts supplemented with E-mix and Ubiquitin. Time-dependent proteolysis was resolved by SDS-PAGE and autoradiography. **(B)** Equivalent assays were performed with non-radioactive IVT products. Target proteins were incubated 1 h in reaction solution containing either extracts or PBS (control). Aliquots of each reaction mix (5 μl) were then loaded directly on a chip via separate channels, each containing dozens of cell units. Target proteins were immobilized to protein chambers via biotinylated anti-GFP antibody. The GFP tag was also used for quantification. GFP signals were calculated from 25-30 cell units per target protein per reaction condition (extracts vs. PBS). Box plots depict ratios of GFP signals (extracts/PBS) at *t*_60 min_. Mean (x) and median (-) are indicated. *\*p* value <0.001. Representative raw data depicting detection on chip of the four target proteins are shown. **(C)** The degradation of Flag-Securin-GFP variant was assayed in tube as described in B. Here, however, aliquots were snap-frozen every 15 min. Time-lapse samples were then loaded on the chip for analysis. Target proteins were immobilized in protein chambers via anti-GFP antibodies and quantified based on GFP fluorescence. Time-dependent degradations of w.t vs. mutant Securin were quantified based on ^35^S signal (standard analysis; *N*=3] and GFP fluorescence (on-chip analysis). Plots depict mean signals and standard error bars. Signals were normalized between max (1, *t*_0_) and min (0, *t*_60 min_) values, allowing proper comparison between two very different methods of detection. **(D)** Equivalent experiment to (C) performed with Flag-Geminin-GFP, except that immobilization was based on anti-Flag antibodies.

Next, we tested whether pDOC can be utilized to obtain reliable kinetic information on protein degradation. Flag-Securin-GFP and Flag-Geminin-GFP were incubated in NDB mitotic extracts for 60 min, and reaction samples were snap frozen in liquid nitrogen every 15 min. After quick thawing, samples representing five time-points were loaded on the chip for signal quantification. Comparable analyses were performed using SDS-PAGE and autoradiography. Non-degradable Flag-SecurinΔ64-GFP and Flag-GemininΔ27-GFP variants were also assayed by autoradiography for control. Considering the vast differences between the two methods of detections, GFP and autoradiography signals were normalized between 0 and 1, meaning that the signal at *t*_60 min_ was subtracted from all other time points. The resulting values were normalized to max signal at *t*_0_ and plotted. As shown in Figure 1C and D, on-chip analysis by pDOC and off-chip analysis by SDS-PAGE-autoradiography exhibited near identical degradation patterns of Flag-Securin-GFP and Flag-Geminin-GFP. Note that reaction cocktails contained 1 μl IVT, 20 μl extracts and 2 μl ubiquitin/E. mix solution, following our standard protocol [18, 27, 34] For each time point, 5 μl reaction mixes were flowed for a period of 3 min through protein chambers with open MITOMI button valves, allowing immobilization of the target protein, while the neck valves were closed. This loading protocol enabled clear visualization of the target protein without the concern of signal saturation (Figure S2). We concluded that pDOC facilitates both end-point and time-course analyses of protein degradation in vitro.

Importantly, Flag-Securin-GFP was immobilized to protein chambers via biotinylated anti-GFP antibodies rather than anti-Flag antibodies. By doing so, we effectively demonstrated that GFP can serve for both immobilization and detection, eliminating the need for two tags. Overall, we find GFP to be an optimal tag for signal detection on-chip. The fusion of a fluorescent protein to a target protein, however, may distort protein folding in ways that effectively limit ubiquitination and proteolysis. In this context, the flexibility of pDOC is particularly advantageous. Figure 3A depicts three configurations by which Securin degradation was analyzed on chip. Securin level was measured at *t*_0_ and *t*_60 min_. Signal detection can be direct, either by GFP or green-Lys. While the former is brighter, the latter allows quantification without tagging. Indirect detection by immunofluorescence was found to be equally informative. Here, short immunodetectable tags minimize the risk of misfolding, but the target proteins must be double tagged because a single tag cannot be used for both immunolabeling and immobilization. Detection by immunofluorescence requires an additional 30 min but we benefit from the bright signal of the fluorophores to which plenty of commercial antibodies are coupled. All three detection protocols revealed the degradation of Securin in NDB mitotic extracts.

**Figure 3:**
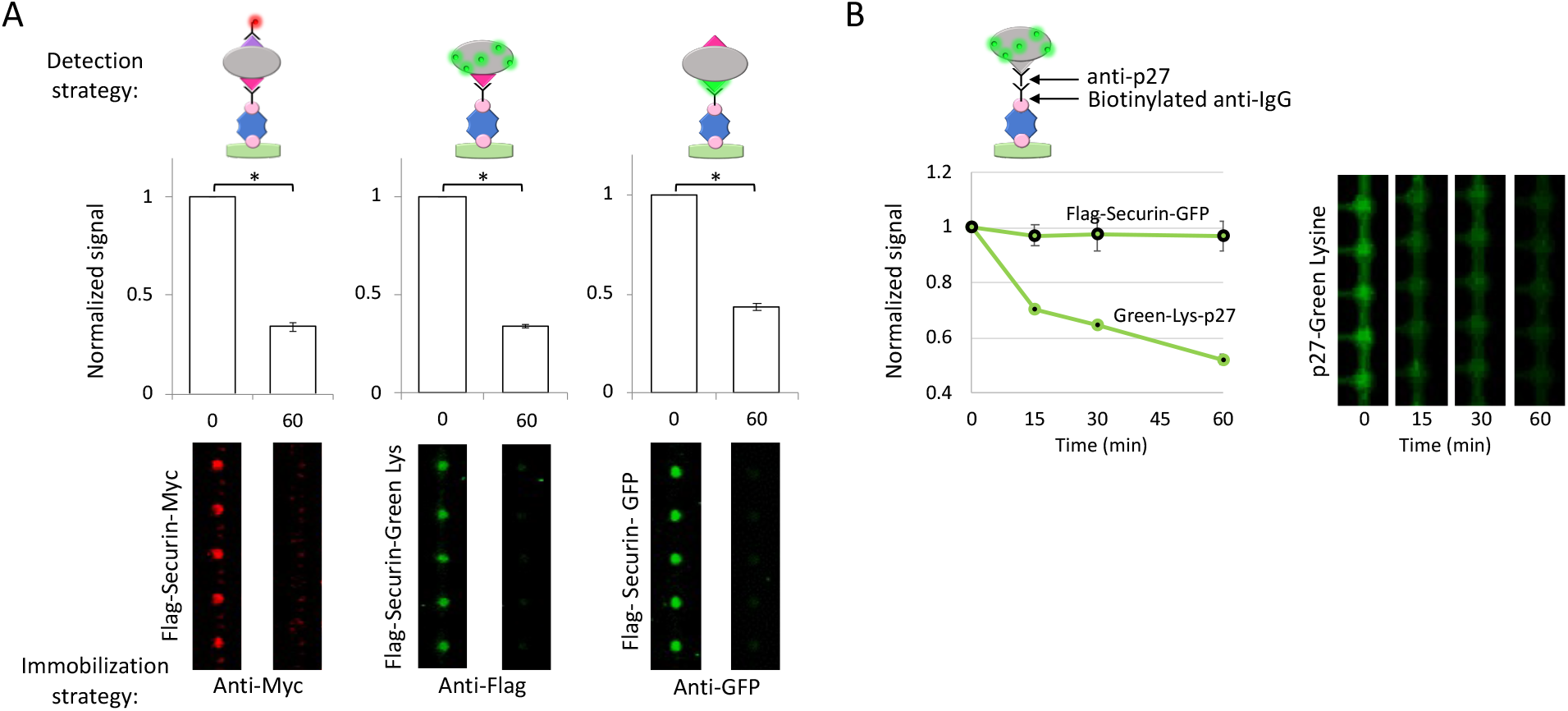
Method versatility. **(A)** Degradation of Flag-Securin-Myc, Flag-Securin-GFP, and Green-Lys-labeled Flag-Securin (IVT products) in NDB mitotic extracts was assayed in tube for 1 h. On-chip analysis was performed in multiple ways: 1) Flag-Securin-Myc was immobilized by anti-Myc antibodies and detected by anti-Flag Cy5-conjugated antibodies; 2) Flag-Securin-GFP was immobilized and detected via the GFP tag; and 3) Green-Lys-labeled Flag-Securin was immobilized via anti-Flag antibodies and detected by the Green-Lys signal. Anti-Myc/Flag/GFP antibodies are biotinylated. Plots and raw data depict protein levels at *t*_0_ vs. *t*_60 min_. Signals are normalized to max values at *t*_0_. Mean values and standard error bars are shown; 20<*N*<30 cell units. \**p* value <0.001. **(B)** Degradation of Green-Lys-labeled p27 (untagged) and Flag-Securin-GFP was assayed in S-phase extracts and analyzed by pDOC. p27 was immobilized via biotinylated anti-mouse IgG and anti-p27 antibodies and detected by Green-Lys. Flag-Securin-GFP was immobilized via biotinylated anti-Flag antibodies. Following incubation with cell extracts, protein levels were quantified by GFP or Green-Lys fluorescence in 15 min intervals. Plots depict mean and standard error values normalized to max signal at *t*_0_. *N*= 25 cell units.

The versatility of pDOC was further demonstrated using tag-free p27 (Figure 3B). Degradation of p27 is mediated by SCF^Skp2^ E3 ligase, rather than APC/C, and is orchestrated with the DNA synthesis (S) phase of the cell cycle [35]. p27 degradation was assayed in extracts from S-phase synchronous HeLa S3 cells and analyzed by pDOC. The protein was immobilized via anti-p27 and biotinylated anti-IgG antibodies, and detected by green-Lys. Analysis by pDOC revealed the stereotypical instability of p27 in S-phase extracts (Figure 3B). The degradation pattern resembled that measured by autoradiography (Figure S3). Thus, pDOC can be utilized for degradation assays of untagged proteins. Furthermore, the method is specific and not restricted to a certain type of cell-free systems.

The advantages of autoradiography-based degradation assays include minimal signal-to-noise ratio, high specificity of the signal and linearity of the assay. However, this does not come without a cost. First, the ^35^S isotope is a short-lived reagent with half-life of ~3 months.

Second, on the gel the signal is spread over a well of 4-5 mm width (standard 10-well gel). Third, in a standard degradation assay, 1-2 μl IVT product is diluted in 20-30 μl cell extracts whose protein concentration is about 20-25 mg/ml. Thus, the amount of IVT loaded per lane is limited by the maximum separation capacity of the gel. De facto, we load 4-5 μl of reaction mix into a well of a standard 10-well mini-gel of 1 mm thick. Fourth, in vitro translation is challenging for large proteins because of ribosome processivity. Thus, although large proteins incorporate more radiolabeled Methionine/Cysteine relative to small proteins, the overall signal of the full-length protein can be impractical for reliable quantification. Fifth, the exposure time for a high-quality signal is typically within a range of 12-24 hrs. Note that the abovementioned points 2-4 are equally relevant when protein degradation is assayed by SDS-PAGE and immunoblotting. As for point 5, western blotting does not require long exposures. Yet, the overall incubation time with antibodies is long. Furthermore, IVT products in immunoblot-based assays must be tagged in order to be distinguished from the endogenous proteins in the extracts. At this juncture, it is important to note that whether degradation is assayed by autoradiography or immunoblotting, informative expression of IVT products must be validated beforehand, in itself, a day long procedure. Equivalent validation by pDOC is instantaneous.

On-chip, the fluorescence signals of Flag-Securin-GFP at *t*_60 min_ were above background level and more noticeable compared to autoradiography (Figures 2A, 3A and S1). Yet, when the signal at *t*_60 min_ was subtracted from all other time-points, the overall degradation patterns of Flag-Securin-GFP, as revealed by pDOC and autoradiography, was similar (Figure 2C and D). This observation suggests that pDOC detects protein residue still lingering in the reaction mix after 60 min incubation, and which are barely detected by autoradiography, if at all. Thus, it can be argued that the sensitivity of pDOC surpasses that of the conventional autoradiography, and if so, pDOC not only facilitates in vitro degradation assays, but also significantly reduces reagent consumption and cost per assay. To test that, we diluted Flag-Securin-GFP in reticulocyte lysate 4- and 10-fold and incubated the substrate in NDB mitotic extracts for 1 hr, while maintaining the original reticulocyte lysate /extract volume ratio of 1/20 (μl). Time point samples were analyzed by pDOC (based on GFP fluorescence). Equivalent experiments performed in parallel with ^35^S-labeled Flag-Securin-GFP, following the conventional assay. To clarify, radiolabeled and non-radiolabeled substrates were expressed simultaneously from the same TNT^®^/DNA solution mix. Furthermore, degradation assays performed a day after delivery of the ^35^S-Met/^35^S-Cys solution to our lab (~1175Ci/mmol), and the gels were exposed to phosphor screen overnight. Yet, whenever the IVT substrate was diluted, the signal obtained by autoradiography was below any acceptable standard, even at *t*_0_, and decreased to barely or undetectable levels after 15 min incubation (Figure 4A). Conversely, analyses by pDOC were informative in all three conditions (Figure 4B). We could detect bona fide signals of 4- and 10-fold diluted Flag-Securin-GFP at *t*_0_ as well as at *t*_60 min_, recording the full dynamics of the protein in NDB mitotic extracts. On a more practical note, we effectively demonstrated that protein degradation can be analyzed with 0.1 μl IVT product and 2 μl cell extracts, thereby saving 90% of the reagents. This feature is particularly valuable in assays which rely on limited biological material, e.g., extracts from primary cells, and normal/pathological tissue samples.

**Figure 4:**
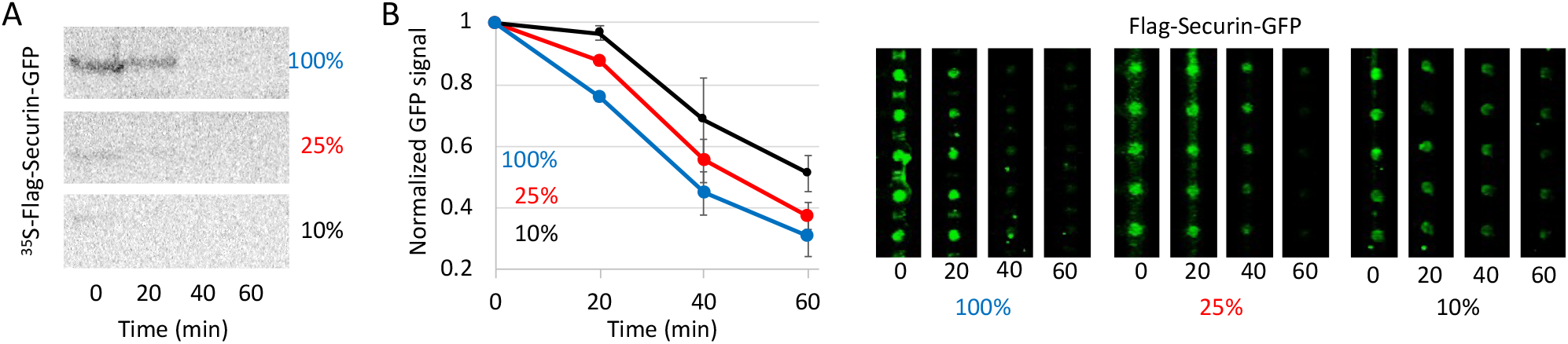
Method sensitivity. **(A)** Time-dependent degradation assay of ^35^S-labeled Flag-Securin-GFP in NDB mitotic extracts. Assays were performed with undiluted IVT product (100%) or following 4x/10x dilution in reticulocyte lysate (25% and 10%, respectively). In all assays, 1 μl substrate was incubated in 20 μl cell extracts. Samples were snap-frozen in 20 min intervals and assayed by SDS-PAGE and autoradiography. **(B)** Equivalent degradation assays performed with nonradioactive Flag-Securin-GFP. Time-point samples were loaded on the chip for immobilization (via anti-GFP-biotinylated antibody) and detection (GFP fluorescence). The plots summarize data from three experiments, 25 cell unites per experiment. Normalized mean and standard error values are shown (left). Representative raw data are shown on the right.

pDOC unveils remanent amount of Flag-Securin-GFP that could not be visualized by conventional methods. Interestingly, while the overall degradation pattern of Flag-Securin-GFP in all three conditions was similar, we noticed a systematic delay in Flag-Securin-GFP degradation as the substrate concentration was reduced. This observation has not been identified previously in our lab, and could be attributed to rate-limiting steps along the process of ubiquitination/degradation of Securin, and perhaps APC/C substrates overall [36, 37].Note that the estimated concentration of Flag-Securin-GFP in the reaction mix was 17 nM (before further dilution in reticulocyte lysate; Figure S4).

### On-chip assay for protein degradation

Until now, we have demonstrated the capacity of pDOC to facilitate analysis of *in tube* protein degradation assays, i.e. the assay takes place prior to its introduction to the chip (Figures 2–4). However, our protocol enables a complete on-chip assay for protein degradation. Reaction samples are loaded on the chip while neck-valves are closed, i.e., without access to chamber I. Target proteins are then immobilized and captured at the protein chamber and remaining materials are washed away (Figure 1A). Technically, the neck valve allows trapping of a target protein in the protein chamber under the MITOMI button valve, and cell-free extracts in chamber I, noted here as ‘extract chamber’ (Figure 5A). The opening of the neck valve enables diffusion of cell extracts into the protein chamber. The motivation for an on-chip assay is threefold: 1) higher-throughput, especially if the target proteins are expressed on-chip, which is possible for MITOMI-based devices; 2) reagent-saving and cost per assay. In fact, 5 μl cell extracts are sufficient to fill a thousand cell units; 3) analysis of protein degradation in real time. The challenge, however, is the limited degradation capacity of <1 nl extracts per cell unit, which never been tested on any platform.

**Figure 5:**
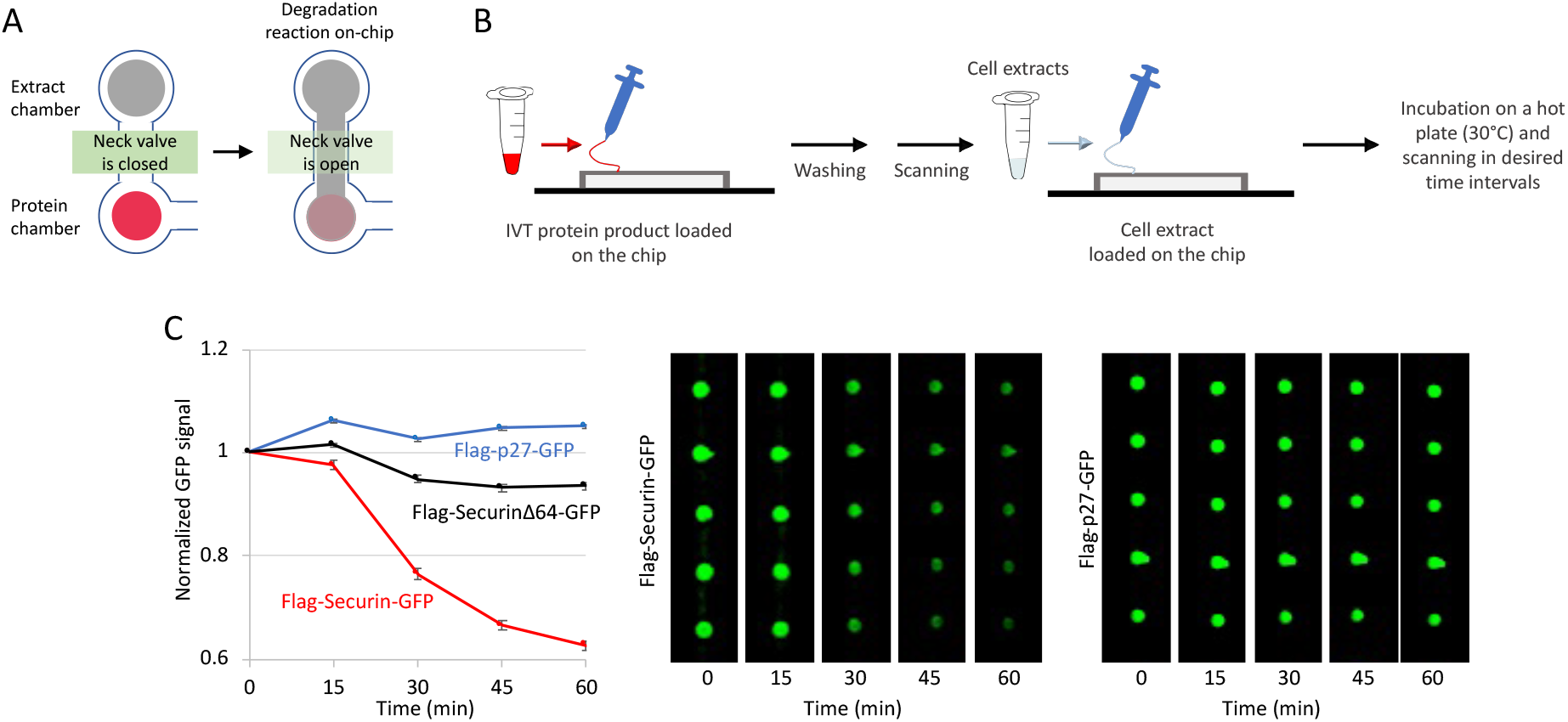
Protein degradation on chip. **(A and B)** Schematic illustration of complete on-chip degradation assay. IVT protein products are immobilized to the surface of the ‘protein chamber’ via biotinylated antibodies (see more information in Figure 1). The closing of the button valve traps the protein. All remains are washed away with PBS. Proper expression and immobilization of the target proteins are validated by scanning. Next, cell-free extracts are loaded into chamber I, i.e., ‘extract chamber’. The reaction begins with the opening of the neck valve and the diffusion of cell extracts into the ‘protein chamber’. The chip is placed on a 30°C hot plate and scanned at desired time intervals to provide kinetic information in real time. **(C)** GFP-tagged Securin and p27 IVT products were immobilized via anti-GFP-biotinylated antibody. Extract chambers were then filled with NDB mitotic extracts that support ubiquitination of Securin, but not of its non-degradable variant (Δ64) and p27 (negative control validating assay specificity). Protein degradation was assayed for 1 hr during which the chip was scanned five times. The plot depicts mean GFP signals normalized to *t*_0_ and standard error bars; *N*=14-25 cell units. Representative raw data for p27 and Securin are shown on the right.

**Figure 6:**
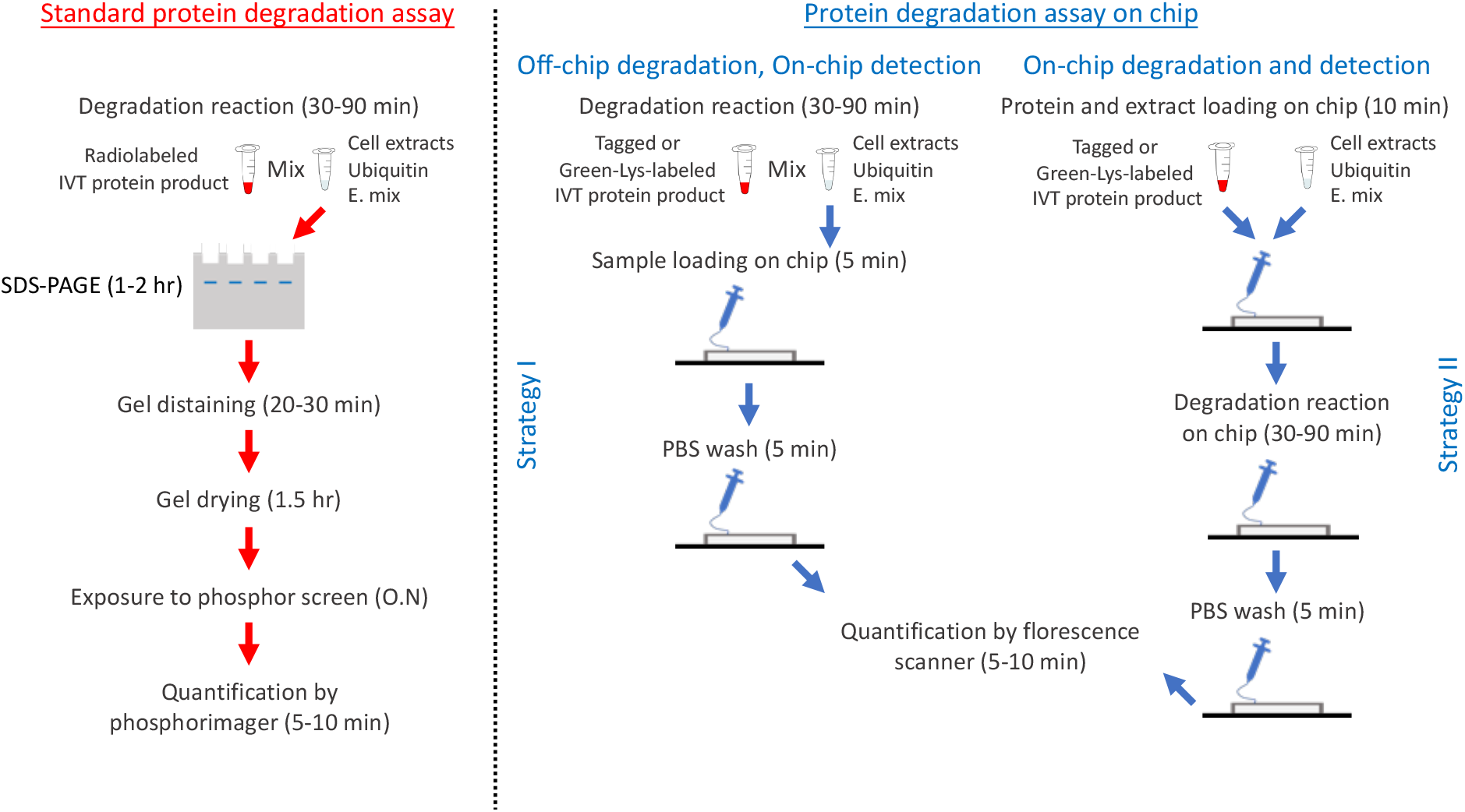
Schematic illustration of standard vs. two on-chip assays for protein degradation. The purpose of all methods is to assay ubiquitin-mediated degradation of in vitro translated (IVT) proteins in cell-free systems comprising native cell extracts, extra ubiquitin and energy-regeneration mixture (E. mix). The standard assay (red pipeline) is typically radioactive. An ^35^S-labeled IVT product (1-2 μl) is mixed with 20-50 μl reaction mixture, and incubated for 30-90 min. Each time point an aliquot (~5 μl) of the reaction mix is harvested. Samples are denatured and resolved by SDS-PAGE. Following drying, the gel is exposed to a phosphor screen. Typically, a satisfactory signal is obtained overnight (O.N). Longer exposures (24-72 hrs) are often needed when the radioactive signal is low due to a variety of reasons. Integrated microfluidics provides two alternative assays, both with significant benefits. Strategy I: The protein degradation is be assayed off-chip. The assay, however, is non-radioactive; the target protein is either fused to a standard tag (e.g., GFP, Flag, Myc, etc.) or translated in the presence of Green-Lys. Time-point samples are either loaded on the chip sequentially or harvested first (boil/snap-freeze) and then loaded simultaneously at the end of the reaction. The target protein is immobilized by tag/protein-specific surface antibodies, all other materials are washed away, and quantification is carried out by one of the strategies detailed in Figure 1B. Strategy II: Here, target proteins and cell extracts are loaded on the chip successively and occupy the protein and extract chambers, respectively. The mixing of the two martials (controlled by the neck valve; Figure 1B) initiates the reaction. Time-lapse scanning provides kinetic information. While Strategy I is simpler, Strategy II is more optimal for high-throughput assays. Both on-chip methods, however, are reagent-saving and considerably shorter than the standard assay.

We decided to test the feasibility of protein degradation on chip. To this end, wt and non-degradable variants of Flag-Securin-GFP as well as Flag-p27-GFP (IVT products) were loaded on the chip and captured in protein chambers along separate channels. All proteins were immobilized via biotinylated anti-GFP antibodies. After washing, NDB mitotic extracts were loaded into the extract chamber and trapped by closing the neck valve. Untrapped materials were washed away. The device was heated to 30°C and scanned to obtain signals of *t*_0_. The opening of the neck valve initiated degradation reactions in all cell units simultaneously (Figure 5A and B), and the chip was scanned in time intervals of 20 min. While the fluorescent signal of non-degradable Securin and p27 remained stable in NDB mitotic extracts throughout the experiment, the signal of Flag-Securin-GFP diminished with time (Figure 5C), revealing the regulated proteolysis of this protein in anaphase.

## Discussion

pDOC is a lab-on-chip platform devised to facilitate and simplify discovery and analysis of protein degradation in physiologically relevant contexts. The chip accommodates hundreds of microchambers in which protein degradation can be assayed promptly and simultaneously using considerably lower quantities of reagents compared to conventional assays. The latter feature is especially crucial for the cell-free extracts whose production can be a bottleneck in terms of time, activity, and amount. Signal detection on pDOC is based on in situ quantification of fluorescent signal. The method is independent of contaminating radioactive materials and yet, with sensitivity that surpasses traditional assays. A comparison between pDOC and the conventional degradation assay is illustrated in Figure 6.

The flexibility of pDOC is significant; the platform facilitates almost all possible experimental designs. First, both tagged and untagged proteins can be assayed, with detection based on incorporation of Green-Lys, fluorescent proteins, or immunodetectable tags. Second, pDOC can be used as an integrated microfluidics column for instant analyses of off-chip degradation reaction. Alternatively, protein degradation can be assayed entirely on-chip and in real time. Third, pDOC allows high-throughput experiments, in which the same protein is tested for degradation in multiple extract/reaction solutions, or the same extracts are applied to dozens of different proteins, simultaneously. Finally and importantly, pDOC is inherently compatible with on-chip in vitro translation [24, 25, 27]. By expressing array of proteins on-chip, one can multiplex the protein targets from tens to thousands, increasing throughput dramatically. The downside of on-chip expression is that the assembly of the device with the DNA microarray is not trivial and currently has to be performed by experts in specialized laboratories.

Ubiquitin-mediated proteolysis is routinely assayed in hundreds of research laboratories worldwide. We devised pDOC to facilitate and simplify in vitro analyses of protein degradation. The method is fast, sensitive, reagent-saving, cost-effective, and inherently optimal for both low- and high-throughput studies. It is also noteworthy that full automation of the platform is foreseeable. We therefore believe that pDOC holds a great potential in basic and translational research.

## Acknowledgements

We thank the Gerber and Tzur lab members for sharing reagents. The Tzur lab is supported by the Israel Science Foundation (ISF) Grant no. 2038/19.

**The authors declare no competing interests**

## Supplementary information

**Figure 1S.**
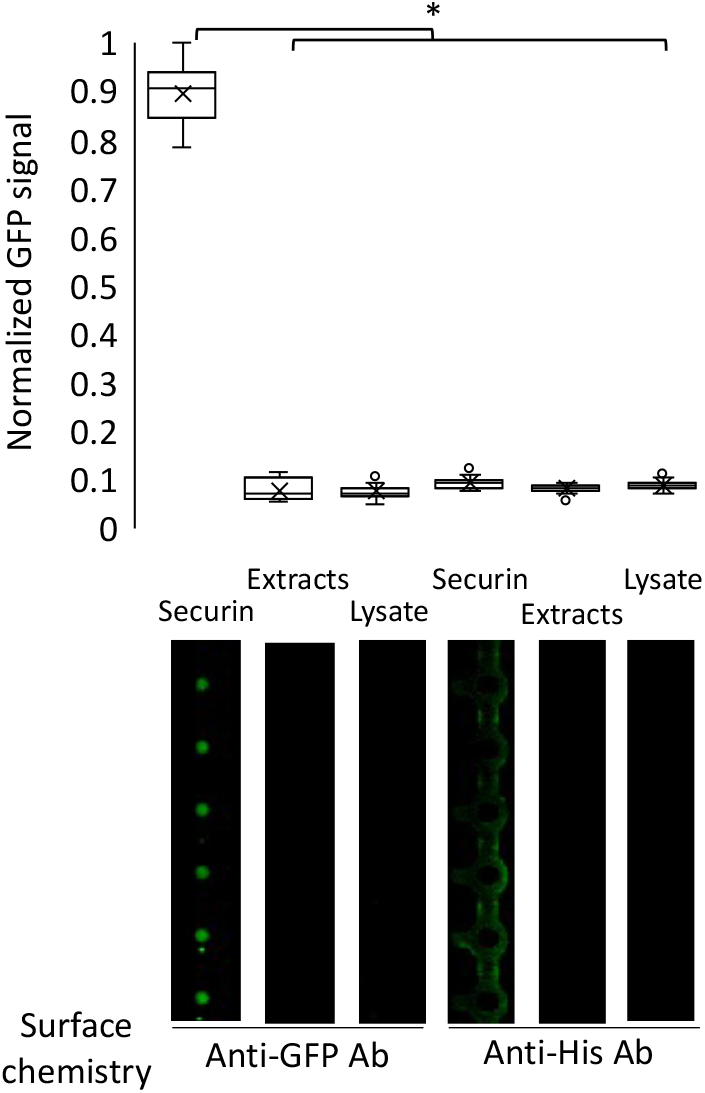
Estimating background and noise signals on chip. An IVT product of Securin-GFP was loaded on a chip, immobilized to protein chambers via biotinylated anti-GFP antibodies, and mixed (on chip) with NDB mitotic extracts (see more information in figures 1 and 6). The GFP signal of the protein was immediately measured. Control experiments performed with 1) NDB mitotic extracts without Securin-GFP (Extracts) to measure the 488 nm-excited autofluorescence of the extracts; and 2) NDB mitotic extracts with reticulocyte lysate to measure the contribution of the lysate to the overall signal of the target protein. A similar set of experiments performed following surface chemistry with biotinylated anti-His antibodies to measure the contribution of non-specific protein interactions to the overall signal coming from the protein chamber. Box plot depicts mean (x) and median (-) signals normalized to maximum level. *N*=39; \**p* value <0.01.

**Figure 2S.**
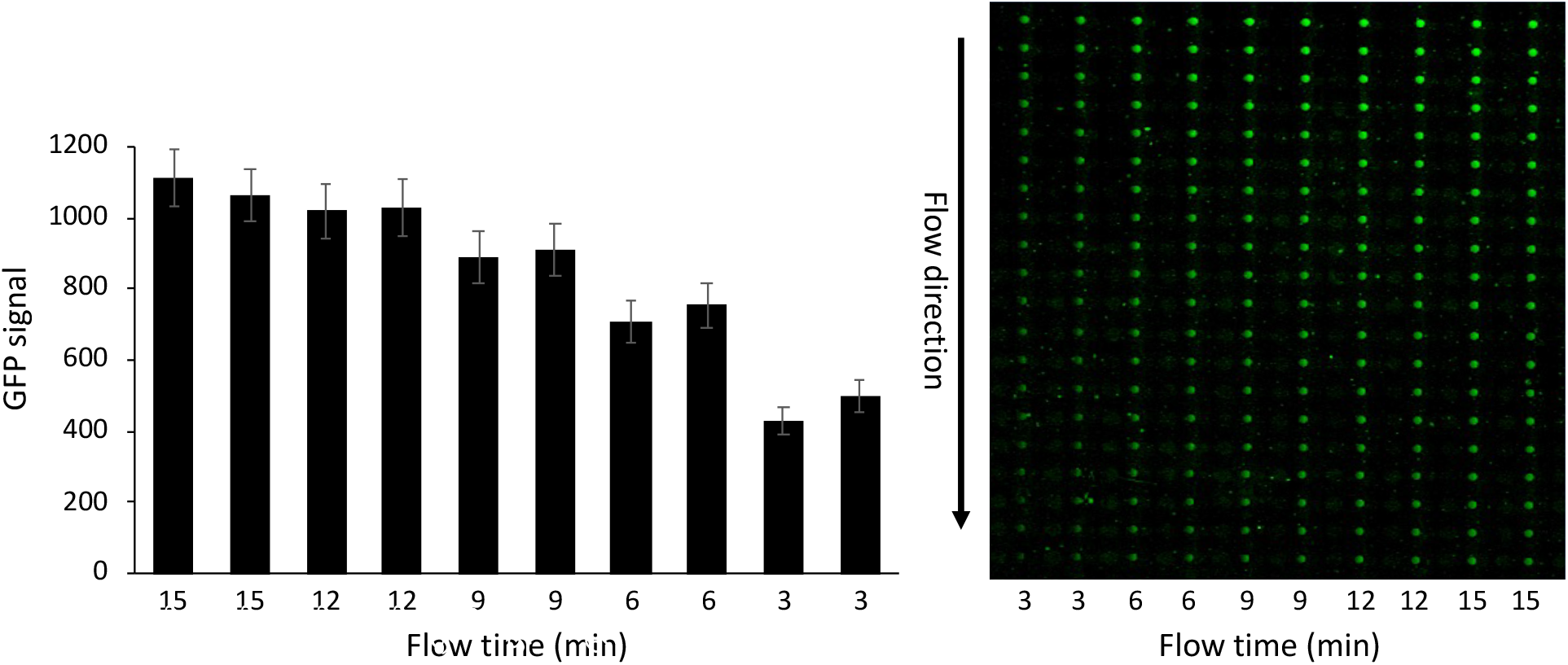
Validating unsaturated signal detection by pDOC. 1 μl IVT product of non-degradable (Δ64) Flag-Securin-GFP was mixed with 20 μl NDB mitotic extracts, following the volume ratio of a standard degradation reaction. Samples were imidiately flown on the pDOC device for various time periods (depeicted) via separate channels (two chanels per sample). Flag-Securin-GFP was immobilized to protein chambers via biotinylated anti-GFP antibodies and detected by GFP fluorescence. The plot depicts mean values calculated from 20 cell units (arbitrary units). Throughout this study, reaction samples were flown for 3-5 min to avoid any risk of signal saturation. To clarify, per experiment the flow time of all samples is identical. Raw data are shown on the right.

**Figure 3S.**
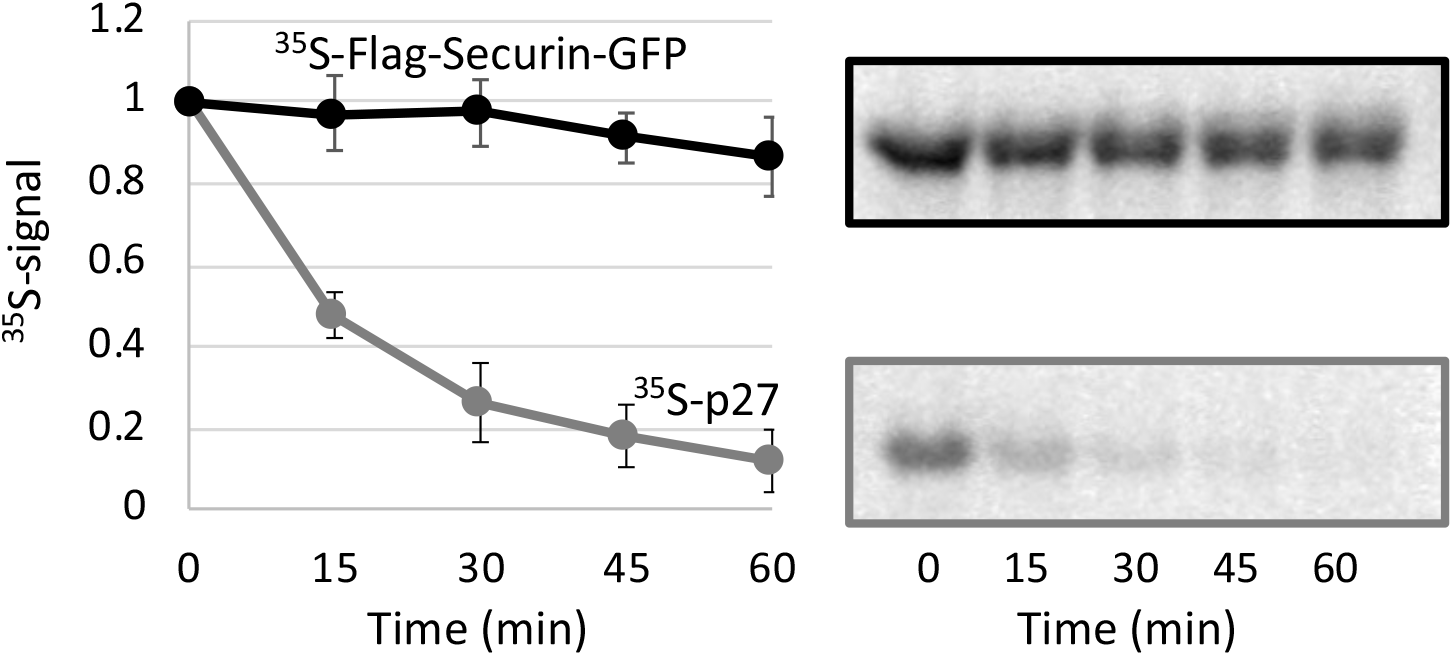
Conventional degradation assay of p27. During DNA synthesis (S) phase, p27 is ubiquitinated by the SCF^Skp2^ E3 complex and degraded whereas Securin remain stable. ^35^S-labeled IVT products of both proteins were incubated in human cell-free system recapitulating S-phase. Protein degradation was assayed by SDS-PAGE and autoradiography (standard protocol). p27, but not Securin, was degraded in this condition. Quantifications and representative raw data are shown. The plot depicts mean and standard error values; *N*=3.

**Figure 4S.**
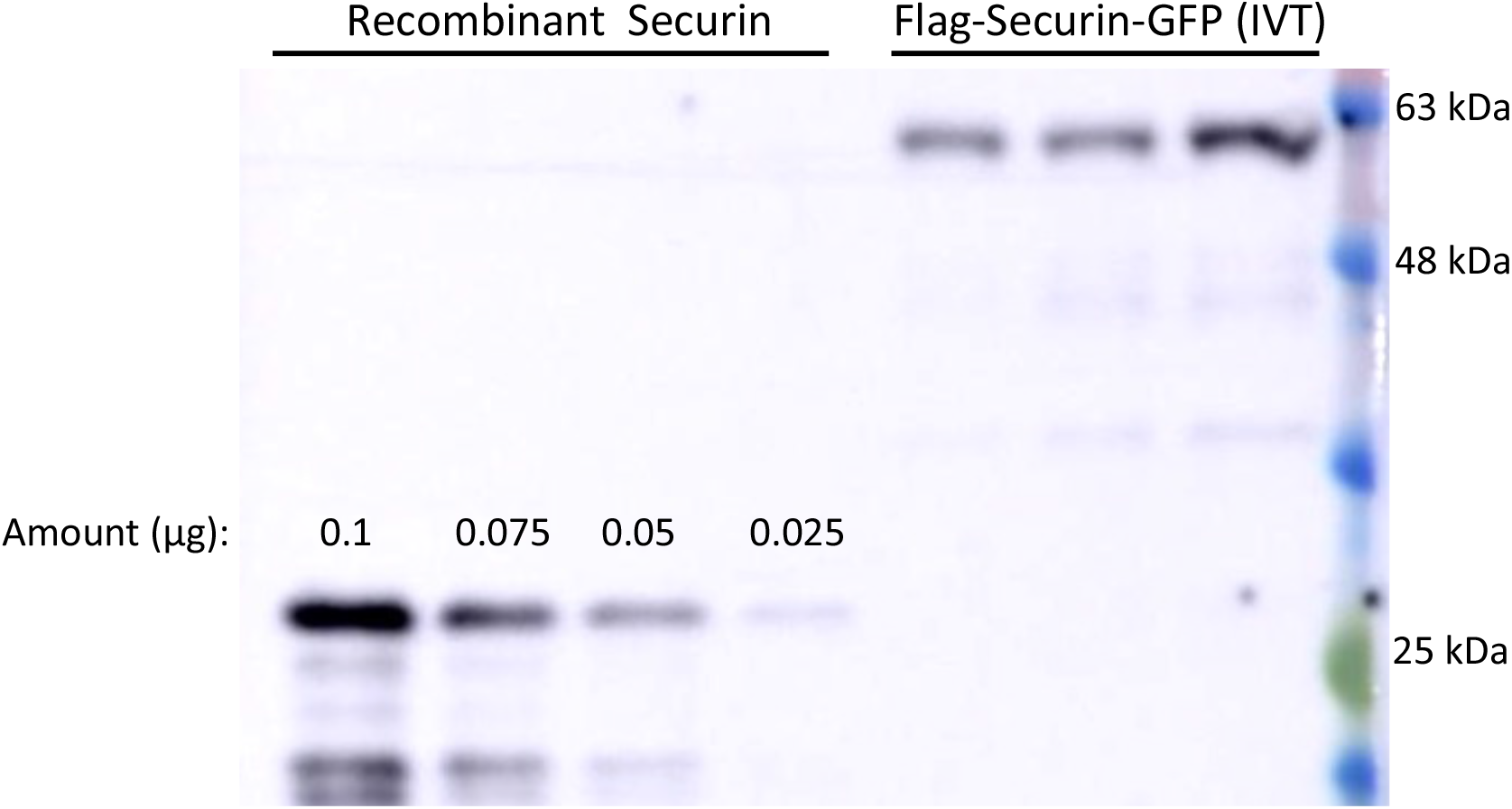
An estimation of Securin concentration in reticulocyte lysate following translation. Various amounts of reombinant His-Securin [34] and 1 μl of three Flag-Securin-GFP IVT products were resolved on SDS-PAGE for Western blot analysis with anti-Securin antibody. The estimated concentration of Flag-Securin-GFP IVT product is 400 nM.

## References

1. Ruz C, Alcantud JL, Montero FV, et al (2020) Proteotoxicity and neurodegenerative diseases. Int. J. Mol. Sci. 21:1–25

2. Ciechanover A (2005) Proteolysis: From the lysosome to ubiquitin and the proteasome. Nat. Rev. Mol. Cell Biol. 6:79–86

3. Emanuele MJ, Enrico TP, Mouery RD, et al (2020) Complex Cartography: Regulation of E2F Transcription Factors by Cyclin F and Ubiquitin. Trends Cell Biol. 0

4. Skaar JR, Pagano M (2009) Control of cell growth by the SCF and APC/C ubiquitin ligases. Curr Opin Cell Biol 21:816–824. https://doi.org/10.1016/j.ceb.2009.08.004

5. Peters JM (1998) SCF and APC: the Yin and Yang of cell cycle regulated proteolysis. Curr Opin Cell Biol 10:759–68

6. Fuchs SY, Spiegelman VS, Kumar KGS (2004) The many faces of β-TrCP E3 ubiquitin ligases: Reflections in the magic mirror of cancer. Oncogene 23:2028–2036

7. Zur A, Brandeis M (2002) Timing of APC/C substrate degradation is determined by fzy/fzr specificity of destruction boxes. EMBO J 21:4500–10. https://doi.org/10.1093/emboj/cdf452

8. Skaar JR, Pagan JK, Pagano M (2013) Mechanisms and function of substrate recruitment by F-box proteins. Nat. Rev. Mol. Cell Biol. 14:369–381

9. Kernan J, Bonacci T, Emanuele MJ (2018) Who guards the guardian? Mechanisms that restrain APC/C during the cell cycle. Biochim. Biophys. Acta – Mol. Cell Res. 1865:1924–1933

10. Choudhury R, Bonacci T, Arceci A, et al (2016) APC/C and SCF(cyclin F) Constitute a Reciprocal Feedback Circuit Controlling S-Phase Entry. Cell Rep 16:3359–3372. https://doi.org/10.1016/j.celrep.2016.08.058

11. Kaiser P, Flick K, Wittenberg C, Reed SI (2000) Regulation of transcription by ubiquitination without proteolysis: Cdc34/SCF(Met30)-mediated inactivation of the transcription factor Met4. Cell 102:303–314. https://doi.org/10.1016/S0092-8674(00)00036-2

12. Bonacci T, Emanuele MJ (2020) Dissenting degradation: Deubiquitinases in cell cycle and cancer. Semin Cancer Biol 67:145–158. https://doi.org/10.1016/j.semcancer.2020.03.008

13. Erales J, Coffino P (2014) Ubiquitin-independent proteasomal degradation. Biochim. Biophys. Acta – Mol. Cell Res. 1843:216–221

14. Glotzer M, Murray AW, Kirschner MW (1991) Cyclin is degraded by the ubiquitin pathway. Nature 349:132–138. https://doi.org/10.1038/349132a0

15. Murray AW, Solomon MJ, Kirschner MW (1989) The role of cyclin synthesis and degradation in the control of maturation promoting factor activity. Nature 339:280–286. https://doi.org/10.1038/339280a0

16. Ayad NG, Rankin S, Ooi D, et al (2005) Identification of ubiquitin ligase substrates by in vitro expression cloning. Methods Enzymol. 399:404–414

17. Nguyen PA, Groen AC, Loose M, et al (2014) Spatial organization of cytokinesis signaling reconstituted in a cell-free system. Science (80-) 346:244–247. https://doi.org/10.1126/science.1256773

18. Wasserman D, Nachum S, Cohen M, et al (2020) Cell cycle oscillators underlying orderly proteolysis of E2F8. Mol Biol Cell mbcE19120725. https://doi.org/10.1091/mbc.E19-12-0725

19. Rape M, Kirschner MW (2004) Autonomous regulation of the anaphase-promoting complex couples mitosis to S-phase entry. Nature 432:588–595. https://doi.org/10.1038/nature03023

20. Yamano H, Gannon J, Mahbubani H, Hunt T (2004) Cell Cycle-Regulated Recognition of the Destruction Box of Cyclin B by the APC/C in Xenopus Egg Extracts. Mol Cell 13:137–147. https://doi.org/10.1016/S1097-2765(03)00480-5

21. Wang W, Wu T, Kirschner MW (2014) The master cell cycle regulator APC-Cdc20 regulates ciliary length and disassembly of the primary cilium. Elife 3:e03083. https://doi.org/10.7554/eLife.03083

22. Khan OM, Almagro J, Nelson JK, et al (2021) Proteasomal degradation of the tumour suppressor FBW7 requires branched ubiquitylation by TRIP12. Nat Commun 12:. https://doi.org/10.1038/s41467-021-22319-5

23. Meyer HJ, Rape M (2014) Enhanced protein degradation by branched ubiquitin chains. Cell 157:910–921. https://doi.org/10.1016/j.cell.2014.03.037

24. Gerber D, Maerkl SJ, Quake SR (2009) An in vitro microfluidic approach to generating protein-interaction networks. Nat Methods 6:71–74. https://doi.org/10.1038/nmeth.1289

25. Glick Y, Ben-Ari Y, Drayman N, et al (2016) Pathogen receptor discovery with a microfluidic human membrane protein array. Proc Natl Acad Sci U S A 113:4344–4349. https://doi.org/10.1073/pnas.1518698113

26. Glick Y, Avrahami D, Michaely E, Gerber D (2012) High-throughput protein expression generator using a microfluidic platform. J Vis Exp. https://doi.org/10.3791/3849

27. Noach-Hirsh M, Nevenzal H, Glick Y, et al (2015) Integrated microfluidics for protein modification discovery. Mol Cell Proteomics 14:2824–2832. https://doi.org/10.1074/mcp.M115.053512

28. Nevenzal H, Noach-Hirsh M, Skornik-Bustan O, et al (2019) A high-throughput integrated microfluidics method enables tyrosine autophosphorylation discovery. Commun Biol 2:1–8. https://doi.org/10.1038/s42003-019-0286-9

29. Ben-Ari Y, Glick Y, Kipper S, et al (2013) Microfluidic large scale integration of viral-host interaction analysis. Lab Chip 13:2202–2209

30. Chen D, Orenstein Y, Golodnitsky R, et al (2016) SELMAP – SELEX affinity landscape MAPping of transcription factor binding sites using integrated microfluidics. Sci Rep 6:. https://doi.org/10.1038/srep33351

31. Wasserman D, Nachum S, Noach-Hirsh M, et al (2021) Elucidating Human Using an Anaphase-Like Cell-Free System. In: Methods in molecular biology (Clifton, N.J.). Methods Mol Biol, pp 143–164

32. Maerkl SJ, Quake SR (2007) A systems approach to measuring the binding energy landscapes of transcription factors. Science (80-) 315:233–237. https://doi.org/10.1126/science.1131007

33. Glick Y, Orenstein Y, Chen D, et al (2015) Integrated microfluidic approach for quantitative high-throughput measurements of transcription factor binding affinities. Nucleic Acids Res 44:51. https://doi.org/10.1093/nar/gkv1327

34. Pe’er T, Lahmi R, Sharaby Y, et al (2013) Gas2l3, a Novel Constriction Site-Associated Protein Whose Regulation Is Mediated by the APC/CCdh1 Complex. PLoS One 8:. https://doi.org/10.1371/journal.pone.0057532

35. Carrano AC, Eytan E, Hershko A, Pagano M (1999) SKP2 is required for ubiquitin-mediated degradation of the CDK inhibitor p27. Nat Cell Biol 1:193–199. https://doi.org/10.1038/12013

36. Meyer HJ, Rape M (2011) Processive ubiquitin chain formation by the anaphase-promoting complex. Semin. Cell Dev. Biol. 22:544–550

37. Lu Y, Lee BH, King RW, et al (2015) Substrate degradation by the proteasome: A single-molecule kinetic analysis. Science (80-) 348:. https://doi.org/10.1126/science.1250834

